# Intranasal oxytocin administration rescues neonatal thermo-sensory deficit in mouse model of Autism

**DOI:** 10.1101/869487

**Authors:** Laura Caccialupi Da Prato, Dina Abdallah, Vanessa Point, Fabienne Schaller, Emilie Pallesi-Pocachard, Aurélie Montheil, Stéphane Canaan, Jean-Luc Gaiarsa, Françoise Muscatelli, Valéry Matarazzo

**Author notes:** corresponding author:; Institut de Neurobiologie de la Méditerranée (INMED), INSERM-Aix Marseille Université, UMR1249, Campus Scientifique de Luminy, 13273 Marseille, France.

## Abstract

Atypical responses to sensory stimuli are considered as a core aspect and early life marker of autism spectrum disorders (ASD). Although recent findings performed in mouse ASD genetic models report sensory deficits, these were explored exclusively during juvenile or adult period. Whether sensory dysfunctions might be present at the early life stage and rescued by therapeutic strategy are fairly uninvestigated. Here we identified that neonatal mice lacking the autism-associated gene *Magel2* fail to react to cool sensory stimuli, while autonomic thermoregulatory function is active. This neonatal deficit was mimicked in control neonates by chemogenetic inactivation of oxytocin neurons. Importantly, intranasal administration of oxytocin was able to rescue the phenotype and brain Erk signaling impairment in mutants. This preclinical study establishes for the first-time early life impairments in thermosensory integration and shows the therapeutic potential benefits of intranasal oxytocin treatment on neonatal atypical sensory reactivity.

## INTRODUCTION

Autism spectrum disorder (ASD) is a developmental disorder characterized by challenges with social interaction, speech and non-verbal communication, as well as repetitive behaviors. However, atypical sensory behaviors are a core aspect of ASD affecting 90% of children (*1*). Importantly, sensory sensitivities have been documented as early as 6 months in ASD infants, preceding considerably the common core features and the diagnosis (*2, 3*). Much of research in animal models of syndromic and non-syndromic forms of ASDs has focused on the social and cognitive difficulties and their underlying mechanisms (*1*). Recent increasing evidences suggest that sensory traits such as tactile, visual, auditory, olfactory, gustatory and heat abnormalities (*4–10*) are present in juvenile/adult ASD models (*11*). However, nor the demonstration of sensory dysfunctions during early post-natal development nor the underlying mechanism have been investigated. These are important steps for early diagnoses and development of therapeutic strategies for ASD.

During the first week of life, sensory integrity is instrumental since neonates have to undertake vital innate behaviors such as nipple-searching and alert calls. Among the various stimuli arising from the external world, sensing any reduction of the ambient temperature is particularly relevant for neonatal pups. Indeed, unlike their homeothermic adult counterparts, neonates are poikilothermic (*12*) and should be kept in close contact with the mother in order to keep their body temperature. In the absence of their warmth-giving mother and being exposed to cool temperatures, neonates generate ultrasonic vocalizations (USV). In fact, during the perinatal period, exposure to low ambient temperatures is considered as a major stimulus for eliciting USV (*13–15*).

The neuronal pathways underlying cool response behavior is still under intensive investigation. At the peripherical level, thermosensory neurons have been described in the skin and also in the Grueneberg ganglion – a cluster of sensory neurons localized at the tip of the nose (*16–19*). It has been proposed that this ganglion influences USV (*20*) generated by rodent neonates to elicit maternal care on exposure to cool temperatures (*14, 15, 21*). Interestingly, neonate mice deleted for the thermoreceptor expressed in these sensory neurons present USV calls impairment after cool exposure (*17*). Following cool exposure, newborns require an effective thermoregulatory adaptative response to produce heat (*22*). During this period, the primary source of heat is produced by the sympathetically mediated metabolism of brown adipose tissue (BAT); also, so called non-shivering thermogenesis (*23*).

Here, using behavioral tests to assess neonatal thermosensory reactivity we discovered the existence of early developmental deficits in thermal sensitivity in neonate mice lacking the autism-associated gene *Magel2*. *MAGEL2* is an imprinted gene highly expressed in the hypothalamus that is paternally expressed and which paternal deletion and mutation cause Prader-Willi (*24*) and Schaaf-Yang (*25*); two syndromes with high prevalence of ASD (27% and 78% respectively). Both syndromes share overlap phenotype including feeding difficulties and hypotonia at birth followed by alterations in social behavior and deficits in cognition over lifespan. The patients have also sensory disorders characterized, in particular, by instability of body temperature manifested by episodes of hyper or hypothermia without infectious causes and which can be fatal in infants (*26, 27*). Moreover, adolescent with ASD present loss of sensory function for thermal perception (*28*).

With the aim to explore the physio-pathophysiological mechanism underlying this thermal deficit we explored peripheral functional activities of both Grueneberg ganglion and BAT. Furthermore, since this oxytocinergic system is considered as a rheostat of adult sensory functions (*29*), a modulator of huddling and thermotaxis behavior in response to cold challenge (*30*) and it is altered in *Magel2*^+/−p^ neonate mice (*31, 32*), we investigated whether central dysfunction of the oxytocinergic system could sustain neonatal thermosensory reactivities and whether OT pharmacological treatment with OT agonists could be a therapeutical approach.

## RESULTS

### ASD-related Magel2 mutation leads to neonatal thermosensory behavior alterations during the first week of life

To assess thermosensitivity in neonates, we developed an experimental procedure based on coolness-induced USV (*17*) (Figure 1A). Wild-type (WT) and *Magel2*^+/−p^ neonates aged from 0 to 6 days old (P0 to P6) were taken from their nests, isolated from the dam, and exposed separately and successively to two different temperatures (ambient: 25°C and cool: 17°C). On analyzing the latency in emitting the first call, which reflects the reactivity of the animals to sense cold, we found that WT neonates presented a latency lower at cool temperatures than under ambient exposure (Figure 1B-E-H-K), while *Magel2*^+/−p^ did not (Figure 1C-F-I-L).

**Figure 1.**
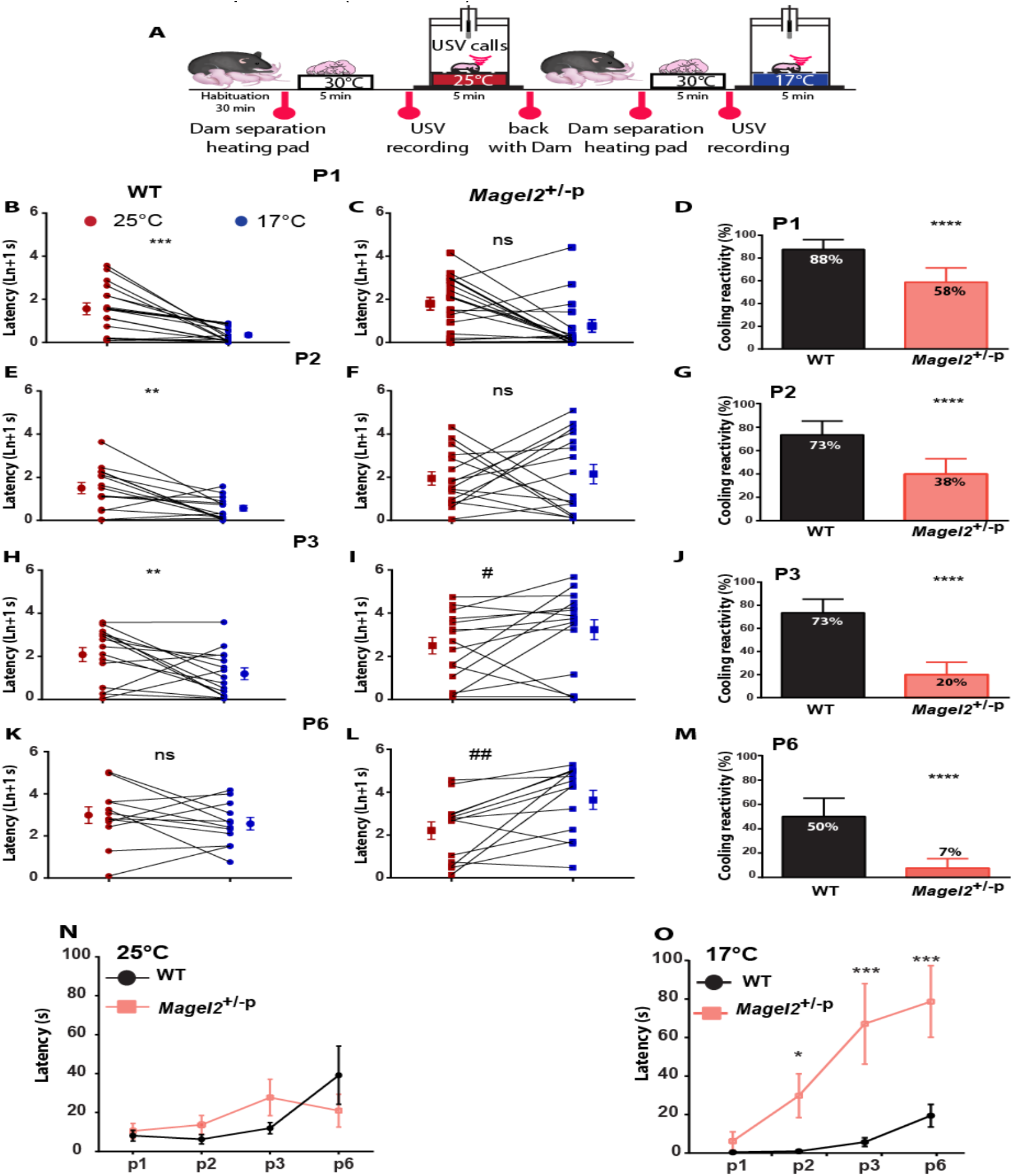
Coolness reactivity failure in Magel2 deficient neonates. **A** : Experimental procedure. After room habituation, neonates are separated from the dam, placed on a heating pad and each neonate is isolated for USVs recording at 25°C for 5 minutes. This procedure is repeated a second time at 17°C exposure and reconducted from P1 to P6 in WT and Magel2+/−p neonates. **B-L**: Before/after graphs illustrating the latency to the first call measured upon exposure at 25°C (red dots) followed by 17°C (blue dots) in WT (B;E;H;K) and in Magel2+/−p (C;F;I;L). WT (25°C vs 17°C): P1: 1.56±0.27 ln+1 s vs 0.34±0.08 ln+1 s; n=16, p=0.0003; P2: 1.48±0.26 ln+1 s vs 0.55±0.13 ln+1 s; n=15, p=0.0054; P3: 2.08±0.32 ln+1 s vs 1.19±0.27 ln+1 s; n=15, p=0.0256; P6: 2.97±0.39 ln+1 s vs 2.56±0.29 ln+1 s, n=12, p=0.3804; Wilcoxon test. Magel2+/−p (25°C vs 17°C): P1: 1.79±0.29 ln+1 s vs 0.76±0.29 ln+1 s; n=17, p=0.0984; P2: 1.95±0.31 ln+1 s vs 2.14±0.45 ln+1 s; n=15, p=0.4037; P3: 2.5±0.38 ln+1 s vs 3.24±0.46 ln+1 s; n=16, p=0.0207; P6: 2.21±0.41 ln+1 s vs 3.65± 0.45 ln+1 s; n=13, p=0.0034; Wilcoxon test. **D;G;J;M**: Bar graphs comparing animals responsive rate of coolness-stimulated USV between WT and Magel2+/−p neonates from P1 to P6. P1: WT: 87.5±8.5 %, n=16 vs Magel2+/−p: 58.82±12.3 %, n=17, p<0.0001; P2: WT: 73.33±11.82 %, n=15 vs Magel2+/−p: 37.5±12.5 %, n=16, p<0.0001; P3: WT: 73.33±11.82 %, n=15 vs Magel2+/−p: 20±10.69 %, n=15, p<0.0001; P6: WT: 50±15.08 %, n=12 vs Magel2+/−p: 7.69±7.69 %, n=13, p<0.0001; Fisher’s exact test. **N-O**: Comparison of the latencies to the first call over the age between WT and Magel2+/−p neonates at 25°C (N) and 17°C (O). Starting at P2, significant age-dependent differences under cool but not ambient exposure were observed: P2: WT 0.96±0.28 s, n=15 vs Magel2+/−p 29.81±11.32 s, n=15; p=0.03; P3: WT 5.71±2.26 s, n=15 vs Magel2+/−p 67.15±20.92 s, n=15, p=0.005; P6: WT 19.47±5.90 s, n=12 vs Magel2+/−p 78.69±18.59 n=12, p=0.007; two-way ANOVA, Bonferroni’s post-test. Data are presented as mean±SEM, *: p<0.05; **: p <0.01; ***: p<0.001; ****: p<0.0001; #: p<0.05; ##: p<0.01; ns: non-significant.

Furthermore, the responsive rate to cool temperature (i.e. the proportion of neonates responsive to cooling) was markedly decreased in *Magel2*^+/−p^ from P1 to P6 compared to the WT (Figure 1D-G-J-M). Comparison of the latencies (Figure 1N-O) between WT and *Magel2*^+/−^ p revealed a significant age-dependent difference under cool but not ambient exposure. This deficit in sensory reactivity was independent of the sex. We also found that dam separation and handling of *Magel2*^+/−p^ did not affect corticosterone levels differently from WT and cannot be linked to this atypical thermosensory reactivity (Supplemental Figure 1).

Furthermore, in order to exclude any potentiation of dam separation we run assays in which USV box temperature was kept at 25°C during the two periods of USV recording (Figure 2A). Using new cohorts of animals, we found that WT performed comparably upon repetitive ambient temperature exposures (Figure 2B-C, and G-H). Moreover, we run experimental paradigm inversion in which another cohort of animals was first exposed to 17°C and then to 25°C (Figure 2D). WT neonates still presented a low-latency response when exposed first to cool *versus* ambient temperatures (Figure 2D, E-F, and G-H). Thus, latency of the first call of isolated neonates was dependent on external temperature. As previously observed, we confirmed that USVs responses to a cool challenge decreased as the pups matured (*33*). At P6, only half of the WT animals were reactive upon cool exposure (Figure 1M). At P8, USV responses were clearly produced independently of the exposed temperature (data not shown). This responsiveness to cool stimuli is independent of mother separation and declines towards adulthood, which is correlated with fur growth, opening of ear canals and the increasing capability of rodents to develop other thermoregulatory capabilities such as seeking comfortable temperature (*13–15, 34*).

**Figure 2.**
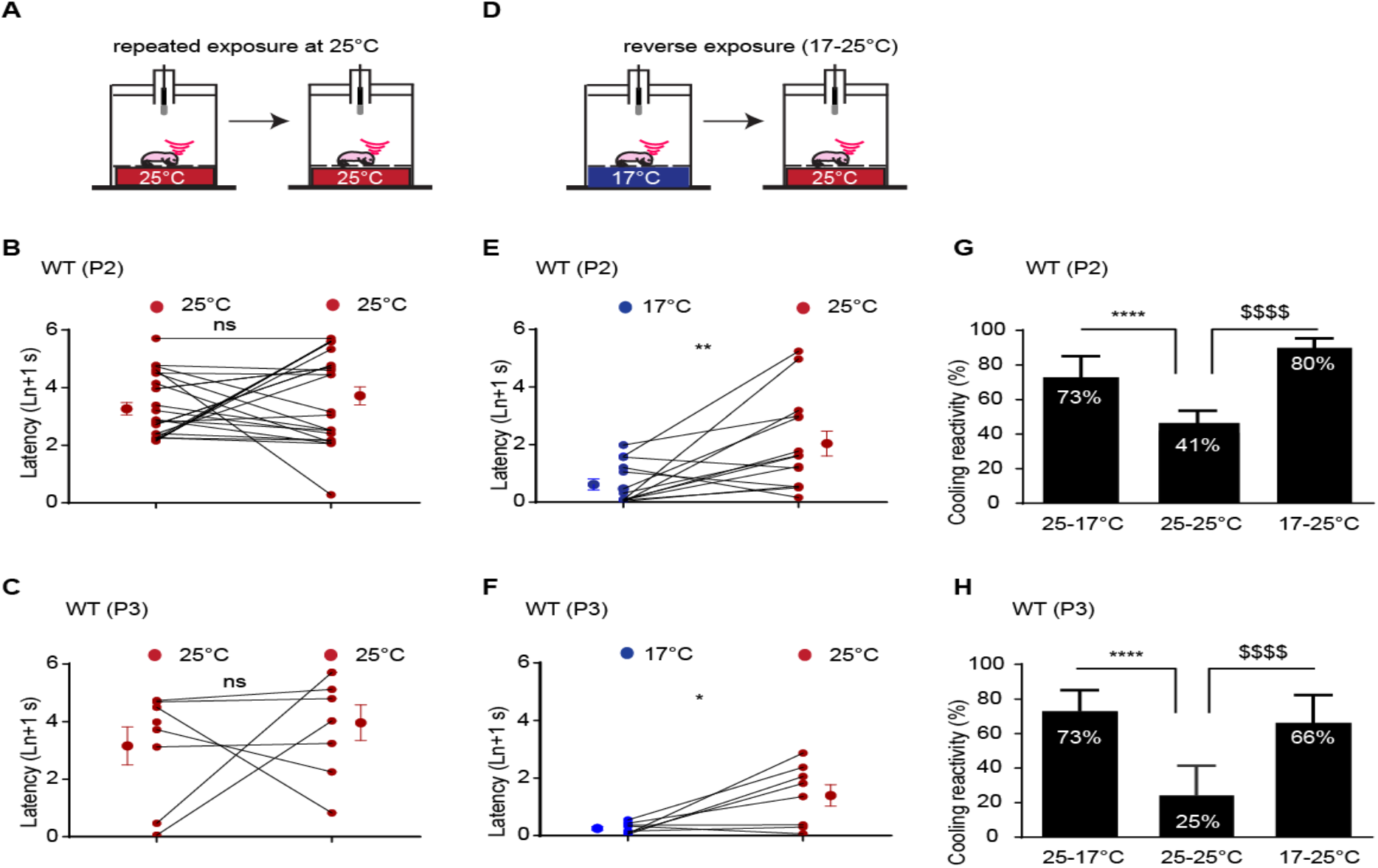
Coolness reactivity in WT upon repeated or reverse temperature exposures. **A-C**: Before/after graphs illustrating the latency to the first call measured upon a repeated exposure at 25°C (red dots) in WT at P2 (B) and P3 (C). **D-F**: Before/after graphs illustrating the latency to the first call measured upon exposure at 17°C (blue dots) followed by 25°C (red dots) in WT at P2 (E) and P3 (F). P2 WT (17°C vs 25°C) : 0.61±0.0.18 ln+1 s vs 2.03±0.43 ln+1 s; n=14, p=0.0.0052; P3 WT: 0.25±0.18 ln+1 s vs 1.40±1.05 ln+1 s; n=8, p=0.039; Wilcoxon test. **G-H**: Bar graphs showing WT animals responsive rate of coolness-stimulated USV at P2 (G) and P3 (H). P2 WT: 25°C-17°C: 73.33±11.82 %, n=15; 25°C-25°C: 41.18±12.30 %, n=17; 17°C-25°C: 85.71±9.70 %, n=14; *p<0.0001; $p<0.0001. P3 WT: 25°C-17°C: 73.33±11.82 %, n=15; 25°C-25°C: 25.00±16.37 %, n=8; 17°C-25°C: 66.67±16.67 %, n=9; *p<0.0001; $p<0.0001; Fisher’s exact test. Data are presented as mean±SEM **: p <0.01; ***: p<0.001; ns: non-significant.

### Cool thermo-sensory behavior impairment in Magel2^+/−p^ neonates is not linked to a deficiency in non-shivering thermogenesis

With the aim to uncover the physio-pathological mechanisms underlying this neonatal thermo-sensory deficit, we first investigate for any thermogenesis dysregulation. Following cool exposure, newborns require an effective thermoregulatory adaptative response to produce heat (*22*). During this period, the primary source of heat is produced by the sympathetically mediated metabolism of brown adipose tissue (BAT); also, so called non-shivering thermogenesis (*23*). Activation of this cold-defensive response is dependent on a thermal afferent neuronal signaling, which involved peripheral thermoreceptors, namely TRPM8, located in dorsal root ganglia sensory neurons of the skin (*35*).

Magel2 being expressed in dorsal root ganglia (*36, 37*), we first analyzed whether peripheral expression of TRPM8 could be affected in *Magel2*^+/−p^ compared to WT neonates. We found no significant difference of the TRPM8 transcript (Figure 3A).

**Figure 3.**
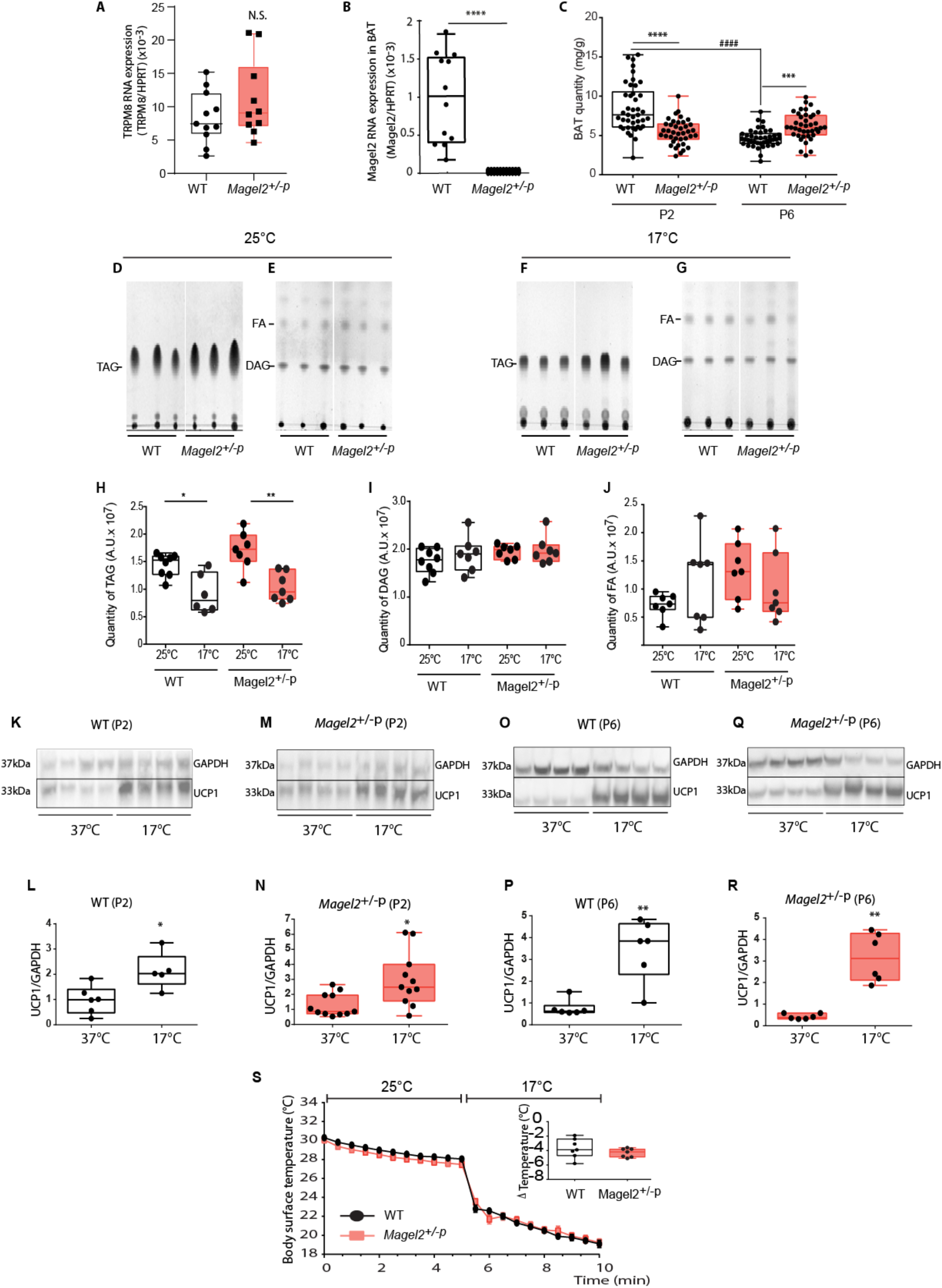
TRPM8 and brown adipose tissue investigations after cool exposure in WT and Magel2^+/−p^. **A**: mRNA expression of TRPM8 in dorsal root ganglia. Quantification of TRPM8 RNA transcripts in Dorsal root ganglia of WT and Magel2+/−p at P2. WT: 0.007 (0.005; 0.012), n=11 vs Magel2+/−p: 0.009 (0.007, 0.016), n=10, p=0.34, Mann Whitney test. Data are presented as median (with interquartile range) (n=10). **B**: Quantification of Magel2 RNA transcripts in WT and Magel2+/−p in BAT and hypothalamus at P2. WT BAT: 1.03×10-3(4.20×10-4; 1.53×10-3), n=12; WT hypothalamus: 3.79×10-2 (3.14×10-2; 4.53×10-2), n=24, *p=0.0006; $p<0.0001; Kruskal-Wallis test, Dunn’s post-test. **C**: Brown adipose tissue (BAT) weight normalized to the body weight of WT and Magel2+/−p at P2 and P6. P2: WT 8.52±0.47 mg/g, n=42 vs Magel2+/−p: 5.25±0.26 mg/g, n=40, p<0.0001; P6: WT: 4.66±0.17 mg/g, n=42 vs Magel2+/−p: 6.18±0.27 mg/g, n=40, p=0.0045, two-way ANOVA, Bonferroni’s post-test; (* between genotype; # intragenotype) **D-J**: Total lipid extraction of WT and Magel2+/−p BAT and thin layer chromatography analysis of TAG, DAG and FA. **H-J**: Quantifications of TAG (H), DAG (I) and FA (J). **K-R**: Immunoblot analyses and quantifications of UCP1 expression after cool exposure at P2 (K-N) in WT and Magel2+/−p (respectively K-L and M-N) and at P6 (O-R) in WT and Magel2+/−p (respectively O -P and Q-R). L: WT P2 :0.99 (0.48, 1.41), n=6, vs 2.03 (1.61, 3.25), n=5, p=0.0173; N: Magel2+/−p P2: 0.86 (0.71, 1.95), n=11, vs 2.49 (1.56, 4), n=11, p=0.0104. P: WT P6: 0.62 (0.57; 0.89), n=6, vs 3.84 (2.31; 4.64), n=6, p=0.0043; R: Magel2+/−p P6: 0.38 (0.32; 0.58), n=6, vs 3.12 (2.11; 4.29), n=6, p=0.0022. **S**: Time course of loss of surface body skin temperature in WT (black line) and Magel2+/−p (red line) of P2 neonates during a temperature challenge (5 minutes at 25°C then 5 minutes at 17°C). Data are presented as mean±SEM. Insert represents the delta loss of surface body temperature before (at 5 min) and after cool exposure (at 10 min) in WT and Magel2+/−p P2 neonates. WT: −3.9 (−4.7; −2.4), n=7 vs Magel2+/−p −4.2 (−4.9;-3.8), n=7, p=0.3648; Mann Whitney test. Mann Whitney test. Data are presented as median (with interquartile range), *: p<0.05; **: p<0.01.

We also found that *Magel2* is expressed in BAT (Figure 3B) and that P2 *Magel2*^+/−p^ neonates had significant decreased interscapular mass BAT compared to aged-matched WT (Figure 3C). As previously shown, we found that in WT the mass of BAT was developmentally down-regulated in WT. Such mass BAT declining in WT was not observed in mutant (Figure 3C).

In order to verify whether this difference of BAT mass could impact non-shivering thermogenesis activity, we investigated BAT function through the analyses of BAT lipolysis and the mitochondrial expression of the uncoupling protein 1 (UCP1) (*38*). In order to observe BAT activation, interscapular BAT tissues from P2 pups were extracted after 1-hour exposure to cool temperature (17°C). Quantitative analyses of BAT lipids, separated by thin layer chromatography (Figure 3D-G), show that although cold exposure induced a significant consumption of triglycerides in both WT and Magel2+/−p pups (Figure 3H), levels of diglycerides and FA remained surprisingly unchanged for both genotypes (Figure 3I-J). Our observation that BAT FA are not consumed upon cold exposure are consistent with recent findings controverting the current view that BAT-derived FA are essential for thermogenesis during acute cold (39).

Next, we found that upon acute cool exposure (17°C, 1h), UCP1 protein expression, a mitochondrial protein activity marker from BAT responsible for non-shivering thermogenesis significantly increases in P2 WT (Figure 3K-L) as well as Magel2+/−p (Figure 3M-N). Similar results were observed at the age of P6 (Figure 3O-P, Q-R). Thus, these results demonstrate that UCP1-mediated non-shivering thermogenesis in BAT is fully active in Magel2 deficient pups. They are also consistent with recent findings showing that UCP1 activation is independent of BAT mass and BAT-derived FA (39). We finally followed skin body temperatures upon cool temperature challenge and found that temperature of Magel2 deficient neonates drop similarly as WT (Figure 3S).

Altogether, our results demonstrate that lack of cool thermosensory call behavior found in Magel2 deficient neonates is not related to their capacity of regulating temperature.

### Cool thermo-sensory behavior impairment in Magel2^+/−p^ neonates is not linked to dysfunction of the peripheral thermosensitive neurons from the Grueneberg ganglion

Peripheral perception to cool temperature is also conducted by the Grueneberg ganglion, a sensory organ located at the tip of the nose (figure 4 A). This ganglion contains sensitive neurons responding to cool temperatures (*18*) and it has been proposed to influence USV (*20*) generated by rodent neonates to elicit maternal care on exposure to cool temperatures (*14, 15, 21*). Interestingly, neonate mice deleted for the thermoreceptor expressed in these sensory neurons present USV calls impairment after cool exposure; a phenotype very similar to what we observe here in *Magel2*^+/−p^ (*17*). We thus ask whether dysfunction of these peripheral thermos-sensory neurons might be affected in *Magel2*^+/−p^. We conducted calcium 2-photon imaging on tissue slices through the Grueneberg ganglion of P2 neonates (Figure 4B-C). Thermo-evoked neuronal activities (obtained by decreasing the temperature of the perfusion solution from 35°C to 17°C) elicited a substantial increase in intracellular Ca^2+^ in both all WT and *Magel2*^+/−p^ animals tested (Figure 4D-E). Thus, the thermosensory neurons of the Grueneberg ganglion are functional in the *Magel2*^+/−p^ neonates.

**Figure 4.**
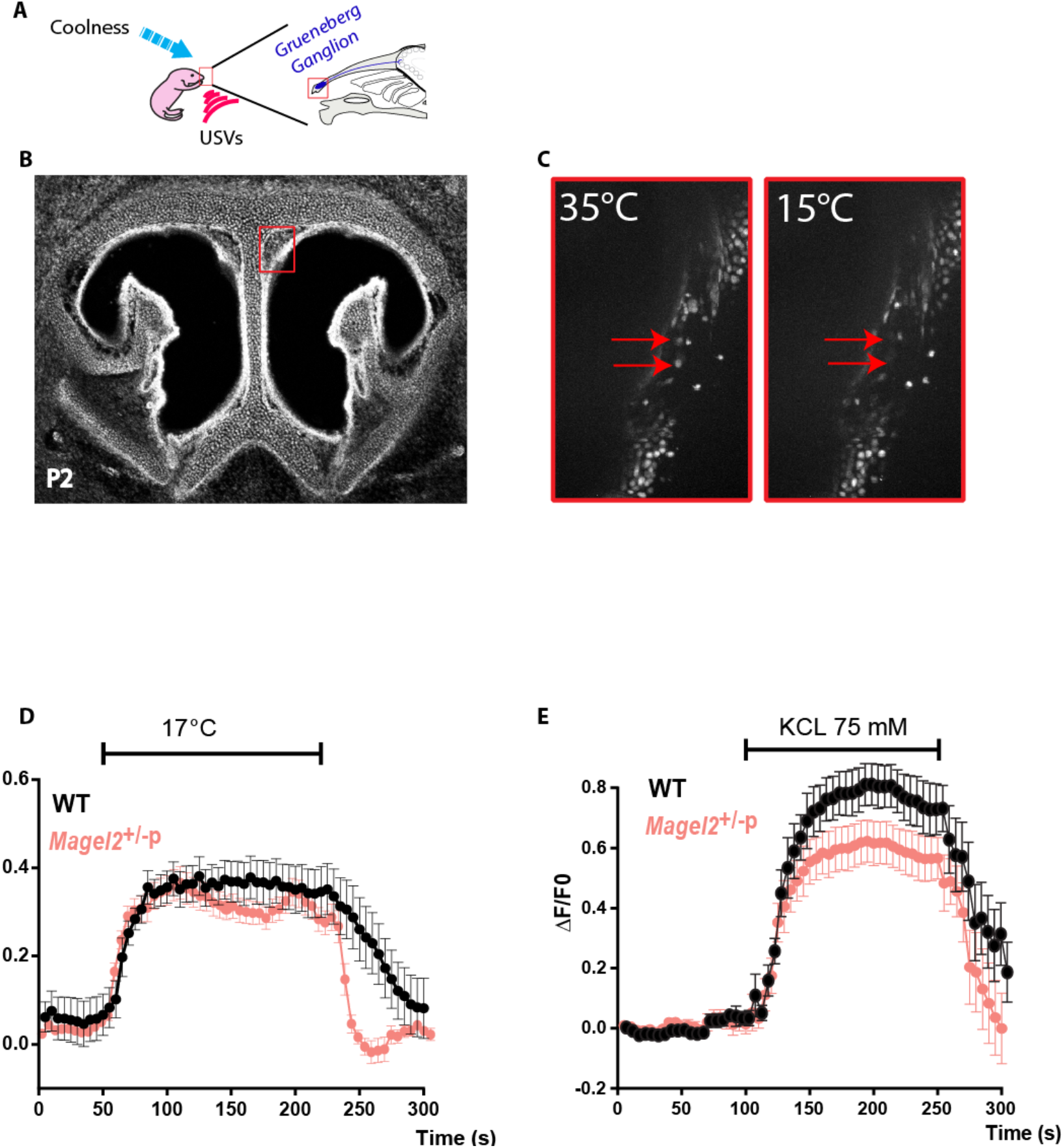
Coolness induced response in the Grueneberg Ganglion (GG) of Magel2+/−p. **A**: Schematic representation of the role and localization of the GG. **B** : Coronal sections of the nasal cavity of a P2 neonate with localization of the GG (red box). **C**: Inserts represent calcium imaging responses after cool exposure. **D**: Representative Ca2+ signals induced by cooling from 35°C to 15°C in WT (n=8) and Magel2+/−p (n=9) GG neurons (respectively black and red). **E**: Representative Ca2+ signals induced in GG neurons by perfusion with KCl (75 mM) were used as a control for viability and responsiveness of tissues slices in WT (n=10) and Magel2+/−p (n=11) GG neurons. ΔF represents change in the ratio of the fluorescence intensity; Data are represented as Mean±SEM.

### Neonatal inactivation of hypothalamic oxytocinergic neurons in WT mimics cool thermo-sensory behavior impairment found in Magel2^+/−p^

Oxytocin (OT) is a main neuropeptide involved in mediating the regulation of adaptive interactions between an individual and his environment (*39*), in major part by modulating sensory systems (*29*). To directly address the involvement of OT neurons in neonatal thermosensory reactivity, we assessed whether inactivation of OT neurons of WT hypothalamus neonates can mimic thermosensory impairment observed in *Magel2*^+/−p^ using DREADD (Designer Receptors Exclusively Activated by Designer Drugs) technology (*40*). This DREADD receptor can be activated by the ligand clozapine N-oxide (CNO) and its metabolite, clozapine; both drugs crossing the BBB (*41*). We restricted hM4Di-mCherry expression to OT neurons by crossing hM4Di-mCherry mice (named here hM4Di) with OT Cre mice in order to drive the expression of the receptor with the OT promoter (Figure 5A-B). These mice were called here OThM4Di.

**Figure 5.**
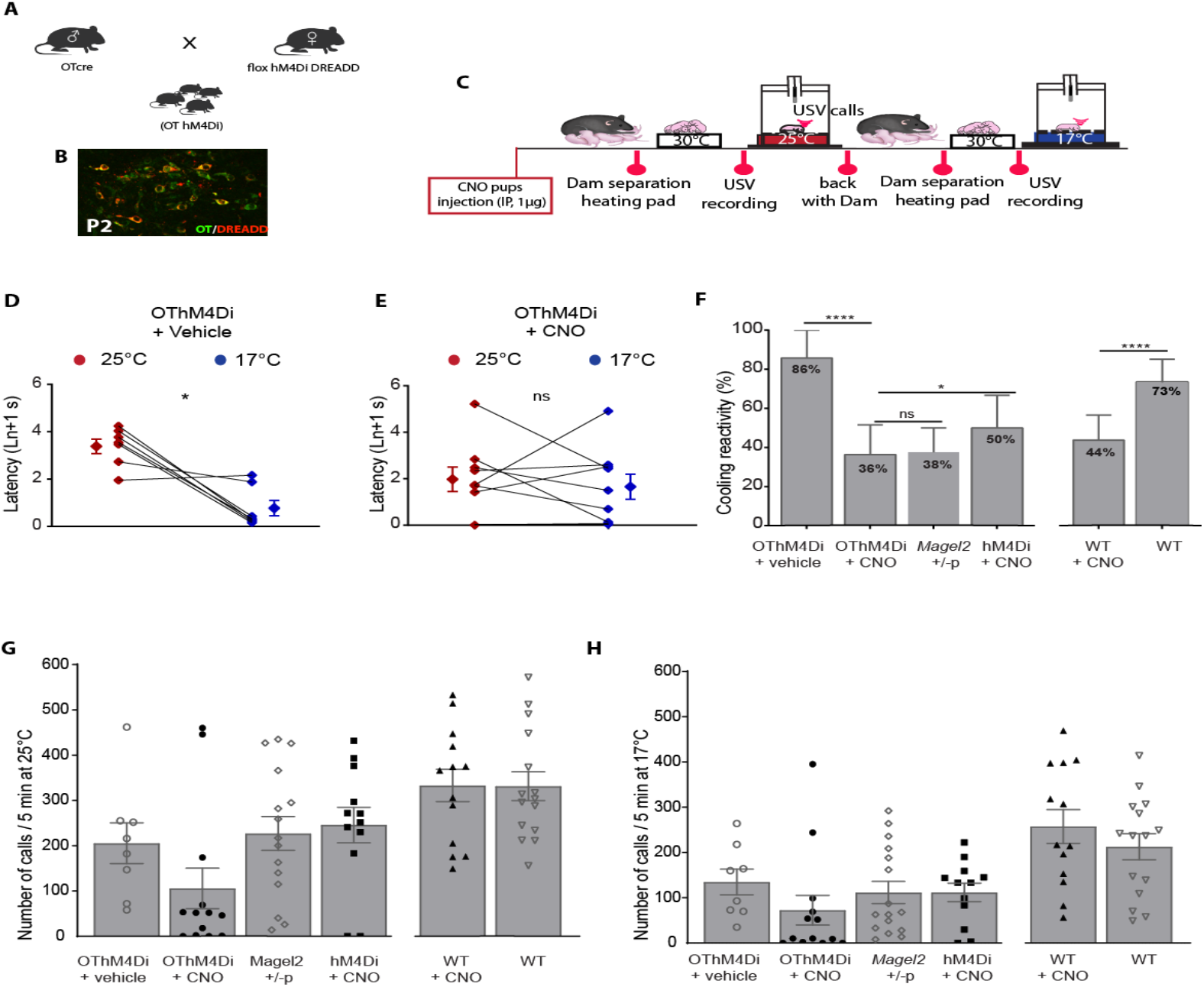
Coolness reactivity failure in WT after oxytocinergic neurons inactivation. **A:** Generation of the hM4Di Dreadd OTCre mice (OThM4Di). **B**: Immunohistochemistry illustrating the expression of hM3Di (red) in OT neurons (green). **C:** Experimental procedure: IP injection of CNO (1 μg) or vehicle was performed in P2 neonates 2 hours before starting experiment. **D-E**: Before/after graphs illustrating the latency to the first call measured upon exposure at 25°C (red dots) followed by 17°C (blue dots) in neonates expressing the hM4Di receptor (OThM4Di) treated with vehicle (D): 3.39±0.3 ln+1 s vs 0.78±0.32 ln+1 s, n=7, p=0.0313 or treated with CNO (E): 1.97±0.52 s vs 1.65±0.54 ln+1 s, n=9, p=0.8203; Wilcoxon test. **F**: Responsive rate of coolness-stimulated USV in OThM4Di neonates treated with vehicle or CNO (85.71±14.29 %, n=7 vs 36.36±15.21 %, n=11; p<0.0001). Cooling reactivity of OT hM4Di neonates treated with CNO was also compared with either Magel2+/−p (37.5±12.5 %, n=16; p=0.1169) or neonates non-expressing the hM4Di receptor (hM4Di) (50±16.67 ln+1 s, n=10, p=0.0315). The last bar graphs illustrate the effect of CNO treatment on WT neonates: WT: 73.33±11.82 %, n=15 vs WT+CNO: 43.75±12.81 %, n=16, p<0.0001; Fischer’s exact test. **G-H**: Total number of calls at 25°C (G) and 17°C (H) in OThM4Di neonates treated or not with CNO and compared with either Magel2+/−p or neonates non-expressing the hM4Di receptor (hM4Di). The last two bar graphs illustrate the effect of CNO treatment on WT neonates. Data are presented as mean±SEM *: p<0.05; **: p<0.01; ***: p<0.001; ****: p<0.0001.

Vehicle or CNO (1μg) was injected into P2 neonates by intraperitoneal (IP) administration two hours before starting thermo-sensory behaviors (Figure 5C). We found that vehicle-treated DREADDs-expressing animals, OThM4Di, presented a significant faster reaction in emitting their first call when exposed to cool *versus* ambient temperatures (Figure 5D); while CNO-treated OThM4Di did not (Figure 5E). Furthermore, the animal responsive rate to cool temperature was markedly decreased in CNO-treated OThM4Di with percentages reaching similar values than *Magel2*^+/−p^ neonates (Figure 5F). CNO treatment did not affect the numbers of USV calls of OThM4Di neonates either at ambient (25°C) or cool temperature (17°C); the number of USV calls being similar to *Magel2*^+/−p^ neonates (Figure 5G-H).

To test for any possible side effects of CNO that were not DREADDs mediated, CNO was administered either to non-DREADDs-expressing (hM4Di) or WT P2 neonates. We found that the responsive rate to cool temperature was damped after CNO administration in WT and in CNO-treated hM4Di neonates (Figure 5F), while the number of USV calls remained similar either at ambient (25°C) or cool temperature (17°C) (Figure 5G and H). However, after CNO treatment responsive rate to cool temperature was still significantly lower in OThM4Di than in hM4Di neonates (Figure 5F). Thus, beside a side effect of CNO which has been reported in other behavioral tests (*42*), our results revealed that *in vivo* inactivation of OT neurons prevents neonates to respond to cool temperature and suggest that OT system can regulate cool sensitivity call behavior in neonates.

### Intranasal injection of Oxytocin rescues cool sensitivity call behavior in Magel2+/−p neonates

We ask whether pharmacological OT treatment could improve thermosensory call behavior in the *Magel2*^+/−p^ during neonatal period (P2). Of the two preferred routes to reach the cerebrospinal fluid, and considering the small size of neonate mice, we found more convenient to administrate OT by intranasal (IN) rather than intravenous route (*43, 44*). New cohorts of neonatal mice were tested for cool thermosensory call behavior with a similar procedure except that neonates received the treatment in between ambient and cool exposures. This procedure allows us to analyze the effect of an acute OT treatment by comparing ambient *versus* cool exposure responses within a same animal (Figure 6A).

**Figure 6.**
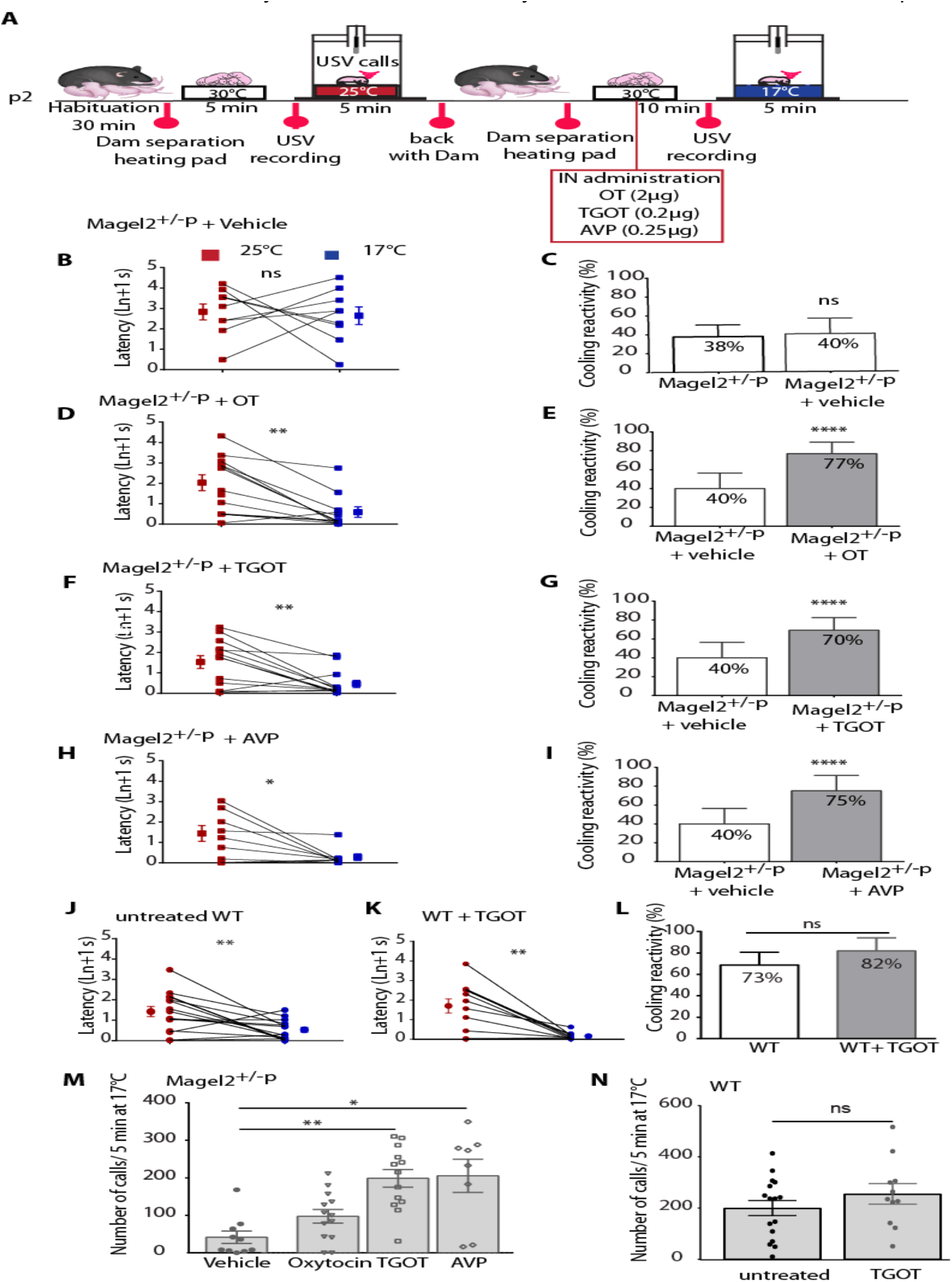
Intranasal oxytocin and oxytocin receptor agonists rescue coolness reactivity in Magel2+/−p. A: Experimental procedure. After room habituation, Magel2+/−p neonates (P2) are separated from the dam, placed on a heating pad and each neonate is isolated for USVs recording at 25°C for 5 min. 10 minutes before repeating the procedure at 17°C, neonates receive an intranasal injection (IN) of Vehicle (NaCl), or Oxytocin (OT, 2 μg) or (Thr4,Gly7)-Oxytocin (TGOT, 0.2 μg) or Vasopressin (AVP, 0.25 μg). B;D;F;H: Before/after graphs represent the latency to the first call measured at 25°C (red dots) and 17°C (blue dots) in Magel2+/−p treated with vehicle (B: 2.83±0.39 ln+1 s vs 2.64±0.43 ln+1 s, n=9, p=0.9102), OT (D: 2.03± 0.39 ln+1 s vs 0.59±0.25 ln+1 s, n=13, p=0.0049), TGOT (F: 1.54±0.31 ln+1 s vs 0.47±0.18 ln+1 s, n=13, p=0.0061) and AVP (H: 1.44±0.39 ln+1 s vs 0.28±0.16 ln+1 s, n=8, p=0.0156); Wilcoxon test. C;E;G;I: Bar graphs showing animals responsive rate of coolness-stimulated USV in Magel2+/−p untreated or treated with vehicle (C: 37.5±12.5 % n=16 vs 40±16.33 %, n=9, p=0.7714), treated with vehicle or OT (E: 40±16.33 %, n=9 vs 76.92±12.16 %, n=13, p<0,0001), or TGOT (G: 69.23±13.32 %, n=13, p<0.0001) or AVP (I: 75±16.37 %, n=8, p<0.0001); Fisher’s exact test. J-K: Latency to the first call measured at 25°C (red dots) and 17°C (blue dots) in untreated (J: 1.48±0.26 ln+1 s vs 0.55±0.13 ln+1 s, n=15, p=0,0054) or TGOT-treated WT (K: 1.71± 0.37 ln+1 s vs 0.16± 0.05 ln+1 s, n=11, p=0.0049); Wilcoxon test. L: Responsive rate of coolness stimulated USV in untreated WT compared with TGOT-treated WT (73.33±11.82 % n=15 vs 81.82±12.20 %, n=11, p=0.2393). M: Total number of calls recorded during 5 minutes in Magel2+/−p treated with vehicle (41.3±16.48, n=10) and compared with OT (97.31±18.16%, n=13, p=0.45), or TGOT (198.50±23.45 %, n=13, p=0.006), or AVP (205.30±44.03 %, n=10, p=0.0035); Kruskal-Wallis test, Dunn’s post-test. N: Total number of calls in untreated WT compared with TGOT-treated WT at 17°C (200.1±29.85, n=16 vs 255.5±39.92, n=11, p=0.5039); Mann Whitney test. Data are presented as mean±SEM, *: p<0.05; **: p<0.01; ***: p<0.001; ****: p<0.0001.

We first verified that handling and IN administration procedures did not affect cool-induced call behavior of *Magel2*^+/−p^ neonates by comparing untreated and vehicle-treated groups. After vehicle treatment (saline solution), *Magel2*^+/−p^ were unable to react to cool exposure since the latency to the first call under cool exposure was similar to ambient exposure (Figure 6B). Furthermore, comparison of the responsive rate to cool temperature (i.e. the proportion of neonates responsive to cooling) under cool exposure revealed insignificant change between these control groups (Figure 6C).

We found that IN administration of OT (2 μg) significantly decreased the latency of the first call of *Magel2*^+/−p^ neonates under cool exposure (Figure 6D) and the responsive rate was markedly increased in *Magel2*^+/−p^ (Figure 6E). Indeed, after OT injection, 77% of *Magel2*^+/−p^ neonates reacted to cool stimuli, a percentage similar to the P2 WT (Figure 6L). Thus, an OT pharmacological treatment is able to rescue the cool thermosensory call behavior deficit of the *Magel2*^+/−p^ neonates.

To better characterize the pathway implicated in the rescue of the cool-induced call behavior, we tested OT agonists. OT and vasopressin (AVP) are closely related nonapeptides that share high sequence and structure homology (*45*). Although one unique receptor exists for OT in mammals, AVP can also bind and activate the OT receptor with the same affinity as OT (*46*). Among the different agonist developed for the OT receptor, [Thr^4^,Gly^7^]OT also referred to as TGOT, has been widely used as a selective OT agonist (*46*); We thus treated *Magel2*^+/−p^ neonates (P2) with either TGOT or AVP and performed USV call recording 10 min after administration of the agonist dose. By analyzing the reactivity of the animals to sense cool temperature, we found that *Magel2*^+/−p^ neonates (P2) presented a significant faster reaction in emitting their first call when exposed to cool *versus* ambient temperature after either TGOT or AVP treatment (Figure 6F and H, respectively). Furthermore, the responsive rate of *Magel2*^+/−^ p neonates to cool temperature was markedly increased in both TGOT and AVP conditions (Figure 6G and I, respectively); reaching similar values as the P2 WT (Figure 6L). We also addressed the action of TGOT in WT neonates and found that it still preserved both the response and the number of USV upon cool temperature exposure (Figure 6J-L and N).

Finally, *Magel2*^+/−p^ pups treated either with AVP or TGOT evoked substantial USV call number upon cool exposure (Figure 6M), with values similar to P2 WT (Figure 6L). Although we cannot completely exclude a minor contribution of the AVP receptors, these data suggest that the rescue of cool thermosensory call behavior is mainly due to activation of the OT receptors.

### Oxytocin rescues Magel2^+/−p^ brain Erk signaling impairment after cool stimuli

Since cool exposure or stress alters the Erk pathways in the brain by reducing Erk activation (*47, 48*) and OT has been shown to block this alteration (*47*), we examined whether this signaling pathways might be altered in *Magel2*^+/−p^ brain neonates. Erk/P-Erk levels were measured from P2 whole brains of WT and *Magel2*^+/−p^ immediately after ambient or cool exposure. Cytoplasmic levels of P-Erk revealed that brain of WT neonates had a significant cool-induced reduction of P-Erk (Figure 7A and C); while *Magel2*^+/−p^ did not (Figure 7B and D). More importantly, IN administration of OT allowed a cool-induced reduction of P-ErK in the *Magel2*^+/−p^ neonates (Figure 7F and H) without affecting the reduction of P-ErK in the brain of WT (Figure 7E and G). Thus, these results highlight a deficit in *Magel2*^+/−p^ brain development and reveal that OT’s ability to reverse cool thermosensory call behavior may act, at least partly, through Erk pathway.

**Figure 7.**
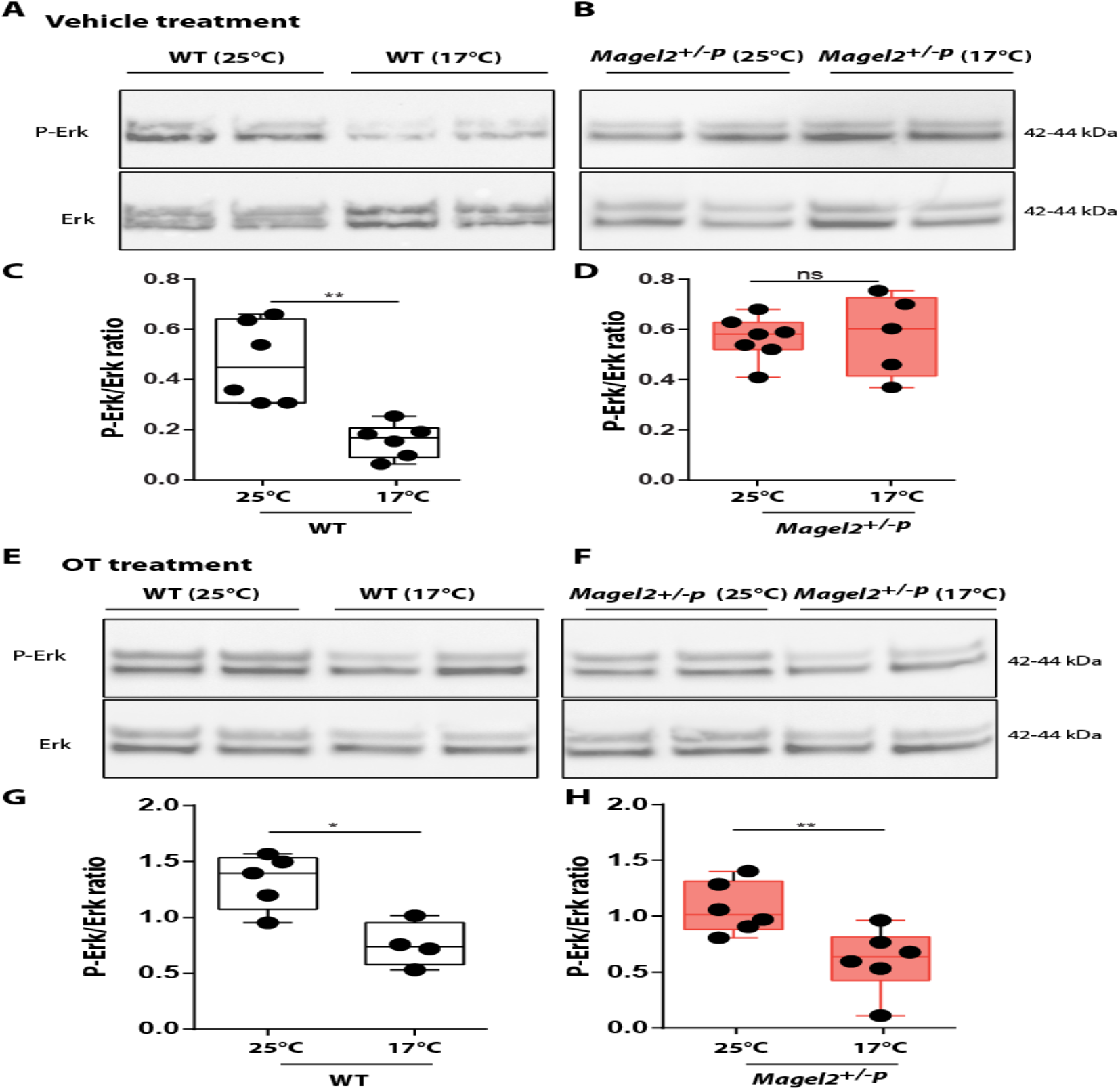
Extracellular signal-regulated kinase (ERK) signaling after cool exposure and oxytocin treatment. A-B: Representative Western blots of cerebral ERK and phosphorylated ERK (pERK) issued from vehicle-treated WT (A) and Magel2+/−p (B) exposed to 25 or 17°C. C-D: Western-blots quantification from WT (C): 0.47±0.07 vs 0.16±0.03, n=6, p=0.0022 or Magel2+/−p (D): 0.56±0.03, n=7 vs 0.58±0.07, n=5, p=0.7424; Mann Whitney test. E-F: Representative Western blots of cerebral ERK and phosphorylated ERK (pERK) issued from intranasal OT-treated WT (E) and Magel2+/−p (F) exposed to 25 or 17°C. G-H: Corresponding western-blots quantification from WT (G): 1.32±0.11, n=5 vs 0.76±0.1, n=4, p=0.0317 or Magel2+/−p (H): 1.08±0.09, n=6 vs 0.62±0.12, n=6, p=0.0087; Mann Whitney test. *: p<0.05; **: p<0.01; ns: non-significant.

## DISCUSSION

ASD research has mainly focused on ASD-related genes and their impact on social and cognitive behavior in adult. However, atypical sensory reactivity that represents early markers of autism and are predictive of social-communication deficits and repetitive behaviors in childhood has been largely overlooked. Although recent findings performed in mouse ASD genetic models report sensory deficits (*4–10*), they were explored during juvenile or adult period. Whether sensory dysfunctions might be present at the early life stage is still unknown. Here we provide the first experimental evidence that newborn harboring deletion in *Magel2*, *a* gene implicated in Prader-Willi and Shaaf-Yang, two syndromes presenting ASD phenotype, exhibit atypical sensory behavior during the first postnatal week.

With the aim to investigate a relevant sensory function during early life, we explored the thermosensory function. Indeed, sensing any reduction of the ambient temperature is particularly vital, since neonates are poikilothermic. In contrast to adult who can adopt diverse strategies in response to cool stimuli such as thermoregulatory behavior and shivering thermogenesis, newborns need to stay with their warmth-giving mother. In absence of this warming, cool exposure elicits an innate behavior characterized by USV emissions. Here we found that deletion of *Magel2* leads to a hyporeactivity in emitting the first call when neonates are isolated from their dam and exposed to cool temperature. This call reactivity deficit is specific to cool exposure and is not the result of an acute dam separation. Moreover, we demonstrate that the oxytocinergic system modulates this neonatal thermosensibility.

By exploring possibilities of peripheral and central origins of this deficit, we found BAT activation upon cool challenge, suggesting that the autonomic neural circuit including expression of thermoreceptors of dorsal root ganglion controlling non-shivering thermogenesis is not affected. Furthermore, functional investigation of the Grueneberg ganglion, a thermosensory system present at the tip of the nose, revealed that this peripheral cool sensor is still active in *Magel2*^+/−p^ neonates. However, we cannot completely exclude a peripheral deficit through a dysfunction of the trigeminal ganglion since it contains thermosensory neurons detecting orofacial cool stimuli and it expresses Magel2 (*49, 50*). The nasal branch of the trigeminal nerve also expresses OT receptors that can be activated after IN administration of OT (*51–53*).

*Magel2*^+/−p^ neonates might encounter not only difficulties in detecting but also in integrating thermo-sensory stimuli. Here, we provided some evidences for the hypothesis of a central origin. First, we found a lack of cool-induced alteration of brain pERK signaling and second, we showed that brain inactivation of OT neurons in WT reproduces atypical thermo-sensory reactivity.

We have provided previous evidences that the oxytocinergic system is altered in *Magel2*^+/−p^ mice and that early OT treatment restores normal motor sucking activity, social and cognitive behaviors in adult mice (*31, 32*). Furthermore clinical trials conducted previously in Prader-Willi babies have demonstrated the efficiency of intranasal oxytocin administration to rescue sucking activity (*54*). Prader-Willi and Schaaf-Yang babies present also sensory disorders characterized in particular by temperature instabilities manifested by episodes of hyper or hypothermia without infectious causes (*26, 27*). Moreover, adolescent with ASD present loss of sensory function for thermal perception (28). Here we demonstrate a new pivotal role of the oxytocinergic system in modulating early life thermosensory function that could be involved in these symptoms.

Although cool-induced cry has been also observed in newborn infants more than 20 years ago (*55*), it is rarely observed nowadays because maintaining the body temperature of the neonate has been emphasized. Measures of early life sensory behavior such as cool-thermosensory call behavior might represent promising avenues for early diagnostic and OT treatment could be considered for therapeutic interventions of this atypical sensory reactivity.

## MATERIAL AND METHODS

### Animals

Mice were handled and cared in accordance with the Guide for the Care and Use of Laboratory Animals (N.R.C., 1996) and the European Communities Council Directive of September 22^th^, 2010 (2010/63/EU, 74). Experimental protocols were approved by the institutional Ethical Committee Guidelines for animal research with the accreditation no. B13-055-19 from the French Ministry of Agriculture. All efforts were made to minimize the number of animals used. 129-Gt(ROSA)26Sor^tm1(CAG-CHRM4*,-mCitrine)Ute^ also known as R26-LSL-hM4Di DREADD were obtained from the Jackson Laboratory (stock #026219) and called here hM4Di DREADD mice for convenience. Due to the parental imprinting of *Magel2* only heterozygous mice (+m/-p) with the mutated allele transferred by the male were used for experiments. The OT-cre mice were obtained from the Jackson Laboratory (stock #24234). In our experiment we used hM4Di DREADD homozygous // heterozygous OT-cre mice (referred here as OT hm4DI).

### USV recording

On the day of testing (P0, P1, P2, P3 and P6), each pup was separated from its littermates and dam after 30 min of habituation to the testing room, placed on a heating pad and each pup were isolated in a box (23 × 28 × 18 cm) located inside an anechoic box (54 × 57 × 41 cm; Couldbourn instruments, PA, USA) for a 5 min test at room temperature (25°C). Then the pup goes back to the dam for 5-10 min and submits a second separation, placed on a heating pad and the USV were recorded during 5 min under cool temperature (17°C). An ultrasound microphone (Avisoft UltraSoundGate condenser microphone capsule CM16/CMPA, Avisoft bioacoustics, Germany) sensitive to frequencies of 10–250 kHz was located in the roof of the isolation box. Recordings were done using Avisoft recorder software (version 4.2) with a sampling rate of 250 kHz in 16 bits format. Data were transferred to SASLab Pro software (version 5.2; Avisoft bioacoustics) and a fast Fourier transformation was conducted (256 FFT-length, 100% frame, Hamming window, and 75%-time window overlap) before the analysis. Recordings were analyzed for the number of calls during the 5 min recording at 25°C and 17°C and for the latency which is the first ultrasound call of the record. The cooling responsive rate was calculated as the proportion of pups responsive to cooling: a pup is responsive if the latency is two time shorter at 17°C than 25°C.

### Animals’ treatment

The solutions injected were isotonic saline (10 μl) for the control mice and 2 μg of OT (Phoenix Pharmaceuticals Inc., cat #051-01) or 0.2 μg (Thr⁴,Gly⁷)-Oxytocin(TGOT) (BACHEM, lot #1062174) or 0.25 μg of Vasopressin (Phoenix Pharmaceuticals Inc., cat #065-07) diluted in isotonic saline (10 μl) for the treated mice. Intranasal administration was performed in P2 mice 10 min before USV recording.

CNO (Clozapine-N-oxide; Sigma-Aldrich, St Louis, MO, USA) was dissolved in dimethyl sulfoxide (DMSO; Sigma-Aldrich, St Louis, MO, USA) and diluted with 0.9% isotonic saline to volume, the DMSO concentrations in the final CNO solutions were 0.5%. 1μg of CNO was administrated by subcutaneous route in a total volume of 10μl. Administration was performed in P2 mice 2 h-2 h30 before USV recording.

### Corticosterone immunoassay

P2 mice were separated from their mother and placed on a heating pad for 5 min., then sacrificed and blood samples were quickly collected. Blood serum was separated by centrifugation (5,000 rpm, 20 min) and stored at −80°C. Serum corticosterone concentrations were measured with corticosterone ELISA kit (Enzo Life Sciences, Farmingdale, NY, USA) according to the manufacturer’s instructions.

### Lipids analysis

BATs from P2 pups were extracted and incubated with 500 μL of Chloroform/Methanol (CHCl_3_:CH_3_OH, 2:1 *V/V*) solution. Organic phase was isolated by centrifugation at 10,000 rpm, washed by 0.2 vol of 0.9% NaCl solution, dried over MgSO_4_ and then concentrated under nitrogen stream.

19 μL of CH_2_Cl_2_ were added per mg of BAT. For TAG analysis, 2 μL of extract containing total lipids were separated on TLC (Silica Gel 60, Merck) by using petroleum ether:diethyl ether (90:10, *v*/*v*) as eluent. For DAG, MAG and FA analysis, 5 μL of the same samples were separated on TLC by using heptane:diethyl ether:formic acid (55:45:1, v/v/v) as eluent. The TLC plates were sprayed with a solution of 5% phosphomolybdic acid in ethanol followed by heating at 120°C in an oven for 5-10 min, to visualize the spots.

Each resolved plate was scanned using a Chemidoc^TM^ MP Imaging System (Bio-Rad), and densitometric analyses were performed using the ImageLab^TM^ software version 5.0 (Bio-Rad) to determine relative TAG content per sample.

### Calcium imaging

Grueneberg ganglion slices (400μm) were incubated with 10 μM of Fura-2-AM (Life technologies) added with Pluronic acid and dissolved in DMSO, for 45 min at 33°C in an oxygenated artificial cerebrospinal fluid (aCSF) dark chamber. aCSF composition was as followed (in mM): 126 NaCl, 3.5 KCl, 2 CaCl_2_, 1.3 MgCl_2_, 1.2 NaH_2_PO_4_, 25 NaHCO_3_ and 11 glucose, pH 7.4 equilibrated with 95% O_2_ and 5% CO_2_. The recording chamber was first filled with warm (30°C) aCSF for 1 min, then perfused with cool (15°C) aCSF for 3 min and then warm with aCSF for 1 min. Images were acquired every 5 sec with an Olympus BX61WI microscope equipped with a multibeam multiphoton pulsed laser scanning system (LaVision BioTecs) as previously described (Crépel et al., 2007). Images were acquired through a CCD camera, which typically resulted in a time resolution of 50–150 ms per frame. Slices were imaged using a 20×, NA 0.95 objective (Olympus). Images were collected by CCD-based imaging system running ImspectorPro software (LaVision Biotec) and analyzed with Fiji software (*56*).

### Protein extraction and Western blotting

Brain and brown adipose tissues were homogenized in RIPA buffer (Thermo Fisher Scientific) with phosphatase and protease inhibitor cocktails (Pierce Protease and Phosphatase Inhibitor Mini Tablets, EDTA-Free) added with 1% Triton (Euromedex, life sciences products) for the brown adipose tissues. Proteins were run on polyacrylamide gel (Bolt 4-12% Bis Tris plus, Invitrogen by Thermo Fisher Scientific) and transferred to a nitrocellulose membrane (GE Healthcare Life Science). Primary antibodies were incubated overnight at 4°C and were as follow: UCP1 (1:1000, Cell Signalling technology, #14670); p44/42 MAPK (1:1000, Cell Signalling technology, #9102); phospho-p44/42 MAPK (1:1000, Cell Signalling technology, #9101); GAPDH (1:1000, Invitrogen#PA1987). Signals were detected using Super Signal West Pico (Thermo Fisher Scientific, #34080) and bands were analyzed with ImageJ.

### Reverse transcription and real time quantitative PCR

Wild-type and mutant newborns were sacrificed at P2 (between 2pm and 4pm). The hypothalamus, BAT and dorsal root ganglia tissues were quickly dissected on ice and rapidly frozen in liquid nitrogen, then stored at −80°C. Total RNA was isolated using the RNeasy^®^ Mini Kit (Qiagen, cat #74104), according to the manufacturer’s protocol and cDNAs were obtained by reverse transcription using QuantiTect^®^ Reverse Transcription Kit (Qiagen, cat #205311), starting with 600 ng of total RNA.

### Temperature

The body surface temperature was measured using an infrared medical thermometer. Temperature’s values were taken every 30 sec during 5 min at room temperature (25°C) and every 30 sec during 5 min at 17°C.

### Statistical analysis

Analyses were performed using non-parametric statistical tools when the size of the samples was small (GraphPad, Prism 6 software) and the level of significance was set at P<0,05. Values are indicated as following: (Q2 (Q1, Q3) or mean±SEM, n, p-value, statistical test) where Q2 is the median, Q1 is the first quartile and Q3 is the third quartile. Appropriate tests were conducted depending on the experiment and are indicated in the figure legends. Mann-Whitney (MW) test was run to compare two unmatched groups and Wilcoxon-Mann-Whitney (WMW) to compare two matched groups. Kruskal-Wallis (KW) followed by a post hoc test Dunn test was run to compare three or more independent groups. Fisher’s exact test was run to compare contingency tables (reactive vs unreactive animals to cool exposure). Two-way ANOVA followed by Bonferroni post-hoc test was performed to compare the effect of two factors on unmatched groups.

## ACKNOWLEDGMENTS

We thank Antonin Vinck and Emmanuelle Brot for their technical help and the members of the animal facility, genotyping and imaging platforms of INMED laboratory. We thank Dr N. Kourdougli, Pr C. Rivera and C Pellegrino for comments. This study has been supported by INSERM, Aix-Marseille Univ., Foundation Lejeune (N°R15117AA) and ANR (PRADOX N°14-CE13-0025), 20 and Prader Willi France foundation grants. L.C was supported by a PhD fellowship from the French Minister for Research and Technology and from the support of A*MIDEX/ANR (Neuro*AMU Neuroschool PhD program) funded by the French Government « Investissements d’Avenir » program. The authors declare no competing financial interests.

---

**Figure 1 supplemental 1.**
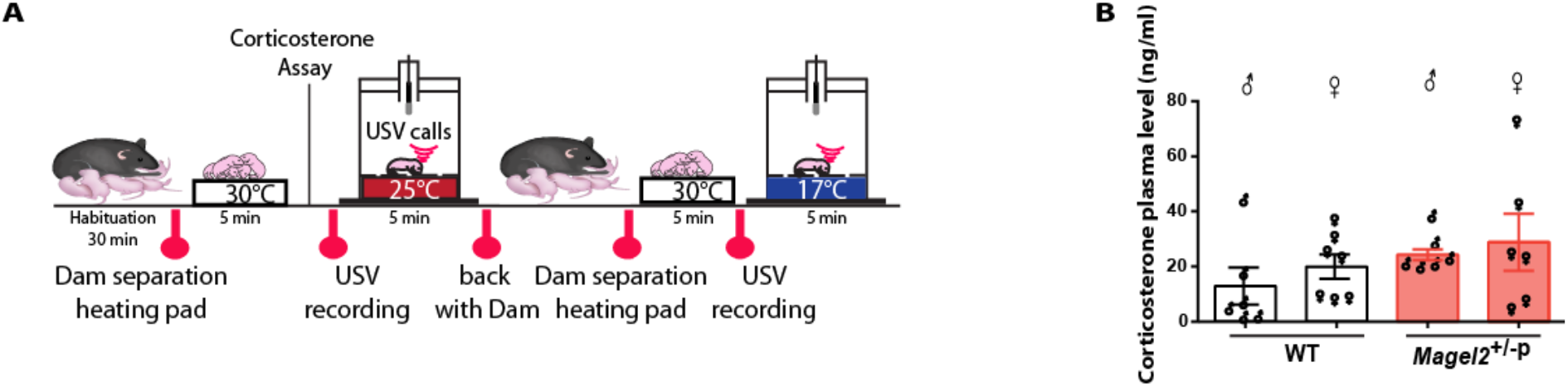
**A**: Experimental procedure. After room habituation, neonates are separated from the dam, placed on a heating pad for 5 minutes and blood samples are collected just before the USV recording. **B**: Comparison between corticosterone plasma levels in female and male of WT and Magel2+/−p measured when neonates are being isolated for USV calls. Males: WT: 12,974±6,711 ng/ml, n=6 vs Magel2+/−p: 24,328±1,971 ng/ml, n=7, p=0.4740. Females: WT 19,983±4,425 ng/ml, n=7 vs Magel2+/−p: 28,831±10,314 ng/ml, n=6, p=0.6723, two-way ANOVA, Bonferroni’s post-test.

